# A *de novo* MS1 feature detector for the Bruker timsTOF Pro

**DOI:** 10.1101/2022.05.02.490258

**Authors:** Daryl Wilding-McBride, Andrew I. Webb

## Abstract

1

Identification of peptides by analysis of data acquired by the two established methods for bottom-up proteomics, DDA and DIA, relies heavily on the fragment spectra. In DDA, peptide features detected in mass spectrometry data are identified by matching their fragment spectra with a peptide database. In DIA, a peptide’s fragment spectra are targeted for extraction and matched with observed spectra. Although fragment ion matching is a central aspect in most peptide identification strategies, the precursor ion in the MS1 data reveals important characteristics as well, including charge state, intensity, monoisotopic m/z, and apex in retention time. Most importantly, the precursor’s mass is essential in determining the potential chemical modification state of the underlying peptide sequence. In the timsTOF, with its additional dimension of collisional cross-section, the data representing the precursor ion also reveals the peptide’s peak in ion mobility. However, the availability of tools to survey precursor ions with a wide range of abundance in timsTOF data across the full mass range is very limited.

Here we present a *de novo* feature detector called three-dimensional intensity descent (3DID). 3DID can detect and extract peptide features down to a configurable intensity level, and finds many more features than several existing tools. 3DID is written in Python and is freely available with an open-source MIT license to facilitate experimentation and further improvement (DOI 10.5281/zenodo.6513126). The dataset used for validation of the algorithm is publicly available (ProteomeXchange identifier PXD030706).

**Author Summary:** In the identification of peptides in mass spectrometry data, much attention has been given to the targeting and extraction of mass spectra produced by fragmentation of precursor ions. However, important information about the peptide is revealed by the data representing the precursor ion itself, such as the peptide’s charge state, mass-to-charge ratio, intensity, and retention time. The timsTOF produces the additional dimension of ion mobility, which provides richer information about the precursor. Although tools exist for the analysis of timsTOF data, they are hampered by limited dynamic range. In this work, we describe a *de novo* feature detector called 3DID that detects peptide features across the full mass range. Our detector can detect more peptides than existing tools across a broader range of abundance, which enables more comprehensive analysis of the data. We believe 3DID will make a valuable contribution to the proteomics toolbox.

## 3 Introduction

For the identification of peptides detected in DDA analysis or extracted in DIA analysis, most peptide identification strategies rely on matching fragment ions with predicted mass spectra from a database of peptides. The precursor ion’s attributes are used to constrain the scope of candidate peptides selected from the database for the matching process (1). The scope is defined by a peptide mass tolerance specified in the search parameters and therefore relies on an accurate determination of the precursor ion’s monoisotopic m/z, charge state, and peak in retention time (2,3). Furthermore, analysis of precursor ions provides much greater depth of analysis on the basis of the stronger signal from precursor ions compared to the intensity of the signal from fragment ions (4). The total mass of the precursor, as well as its position in retention time and mobility, encodes information that is not obtainable solely from analysis of fragment ions.

The timsTOF provides an extra degree of separation of convoluted peptides through their collisional cross-section (5), though resolving peptide features in its MS1 data has formidable challenges. The factors that make challenging the finding and resolving of peptide features in four dimensions include peptide features appearing in the presence of noise that can mimic signal, and feature overlap in m/z, mobility, and retention time as peptides coelute in the liquid chromatography. Although there is variability in the size and shape of features, their typical dimensions are 2 ± 0.3 Th wide in m/z, 44 ± 14 TOF scans through mobility, and 6.6 ± 2.4 seconds in retention time. In a complex sample such as HeLa, there may be 100,000 features in the raw data from the instrument.

An example of candidate features observed in a single frame of MS1 timsTOF raw data is shown in Figure 1. The x-axis is the m/z dimension, and the y-axis is the CCS dimension. The retention time dimension is the series of subsequent frames through time. The frame rate depends on the instrument configuration; in this example the MS1 frame period was 500 milliseconds.

**Figure 1.**
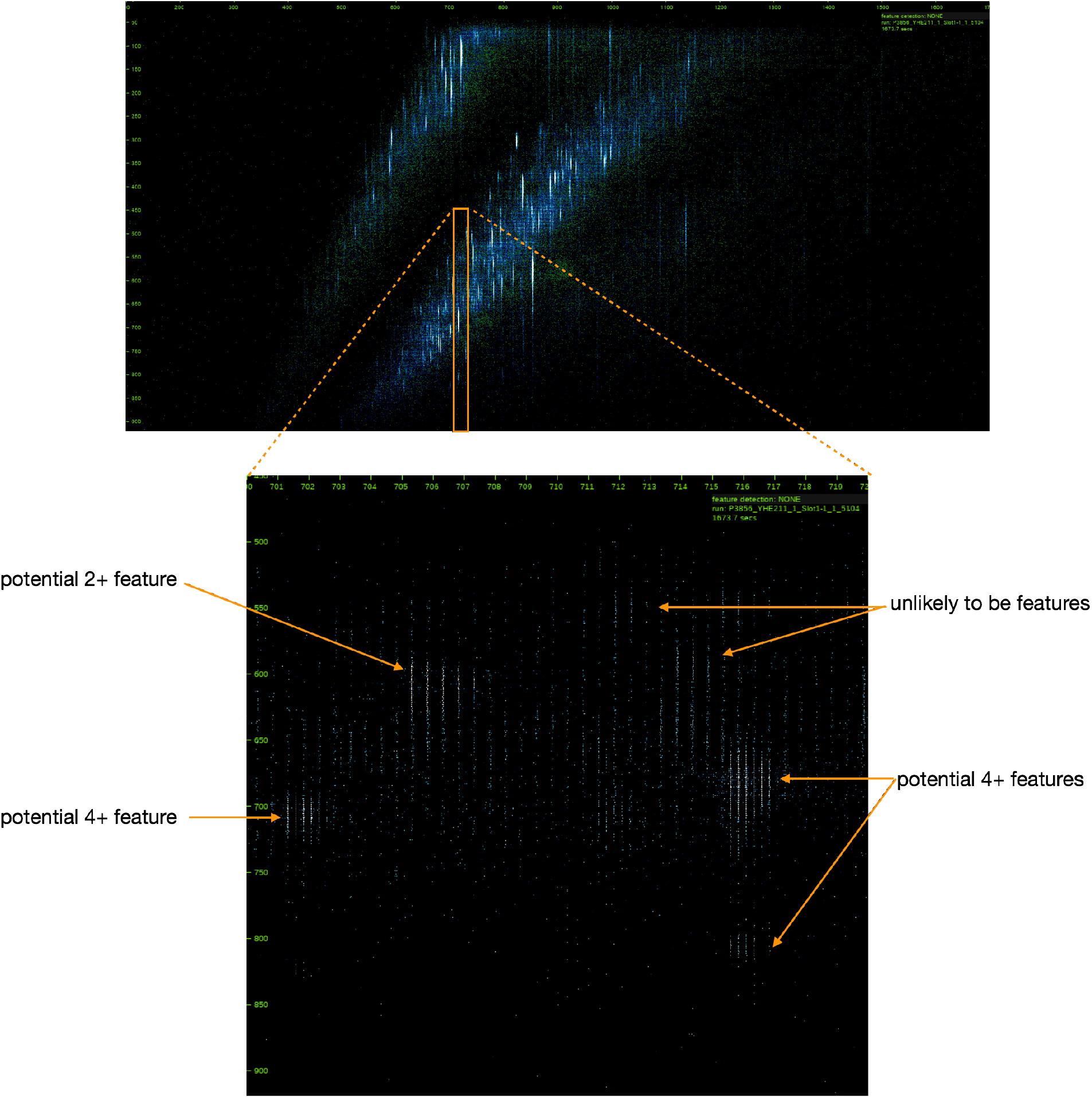
the lower panel shows a small region (20 Da wide) of a raw data MS1 frame shown in full in the upper panel; some potential peptide features are indicated

**Figure 2.**
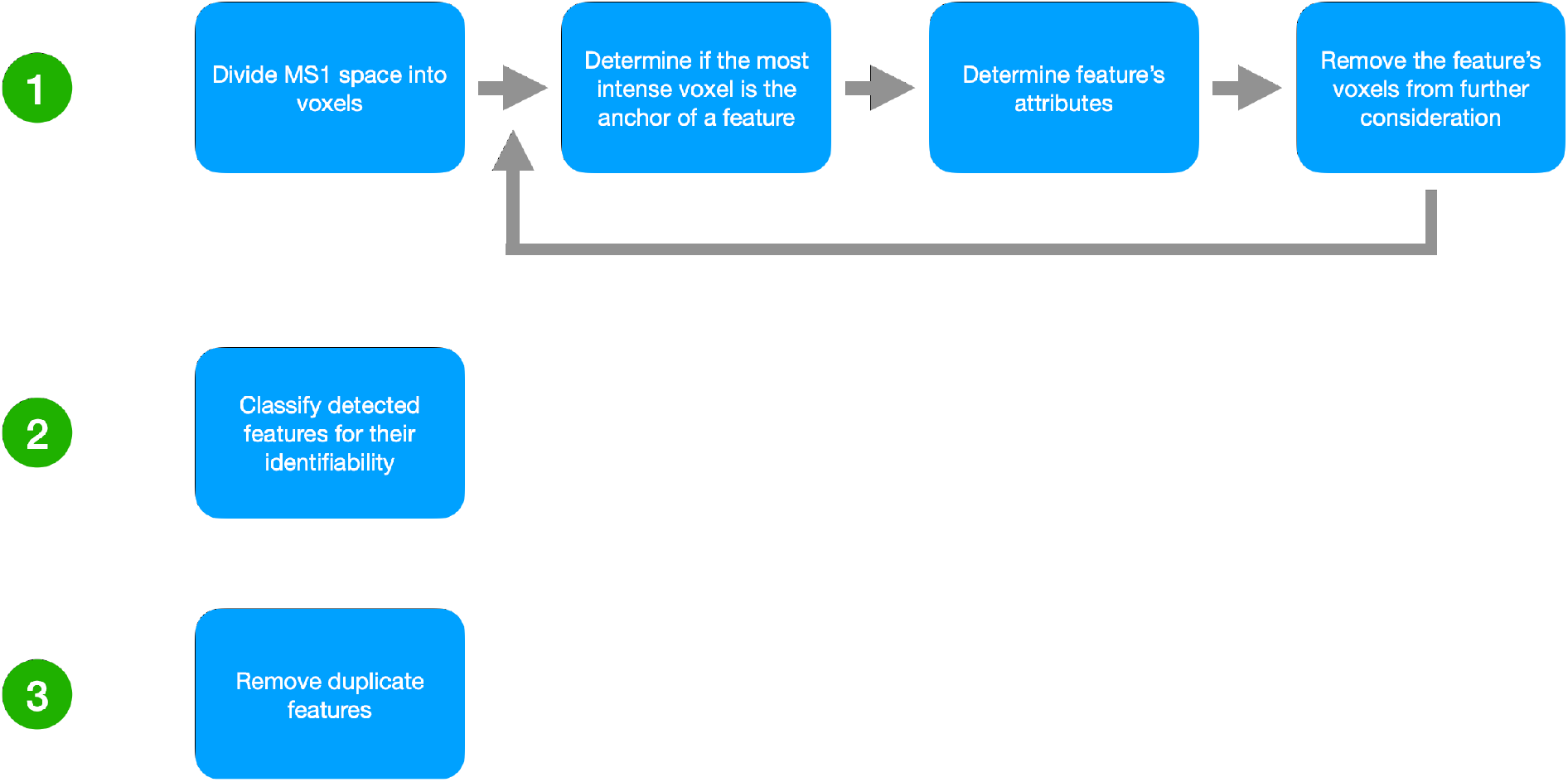
3DID workflow

Previous examples of software that detect peptide features in timsTOF MS1 data across the full mass range includes MaxQuant (6) and Biosaur (7). The approach used by MaxQuant is to average the raw intensities with a Gaussian kernel to form a cube of data with axes of ion mobility index, retention time, and m/z. The cubes are sliced in the mobility dimension, yielding a map of signal intensity as a function of m/z and retention time. Every Nth slice (where N is user-configurable and defaults to 3) is processed without considering mobility with MaxQuant’s standard approach to segment the base of each peak in each m/z-RT plane. The base areas of peaks are then clustered between consecutive planes to form the feature in 4D. MaxQuant performs *de novo* feature detection in MS1 but only reports the features that were included in the isolation regions selected for fragmentation; these are recorded in the APL tables for input to its search engine Andromeda (8).

Biosaur extends the three-dimensional approach of Dinosaur (9) to include the CCS dimension. Raw intensity readings are grouped into ‘hills’ in retention time and CCS within a m/z tolerance. Readings are absorbed into existing hills if their m/z is within tolerance, otherwise a new hill is formed. Isotopic peaks are formed in 4D by gathering readings in consecutive frames and mobility scans, with user-definable tolerance for missed frames or scans for a reading to be considered part of an existing hill.

Here we present a *de novo* MS1 feature detector for the Bruker timsTOF, called 3DID (three-dimensional intensity descent), that performs a comprehensive survey of precursor ions, and detects many more qualified peptide features than MaxQuant and Biosaur.

## 4 Results

Searching for peptide features in vast, sparse, noisy data in a performant manner is an extremely challenging pattern recognition problem. 3DID surveys the entire MS1 data space for the characteristic signature of a peptide’s precursor ion. It does this without prior knowledge of where they might exist. Upon detection of a feature, 3DID determines its attributes: monoisotopic m/z, charge state, and the apex of the feature in mobility and retention time. It also performs quality checks such as the alignment of isotopic peaks in the mobility and retention time dimensions.

3DID leverages knowledge of the predictable structural pattern of peptide features (Figure 3). The first characteristic of the pattern is that peptide features appear as a series of isotopic peaks, each peak resulting from probabilistic combinations of isotopes in the peptide molecule’s composition (10). Second, each peak is comprised of a variable number of intensity readings that present in three dimensions: m/z, CCS, and retention time. The observation of multiple readings with intensity greater than the average intensity of other readings in their vicinity is an indication of a base peak. Third, isotopic peaks have a known width in m/z, determined by the resolution of the instrument. They are Gaussian in the m/z, mobility, and retention time dimensions (11). Finally, isotopic peaks in a series are a known distance apart in the m/z dimension (12), the distance between them being determined by their integer charge state. Isotopic peaks in a series are aligned in the mobility and retention time dimensions.

**Figure 3.**
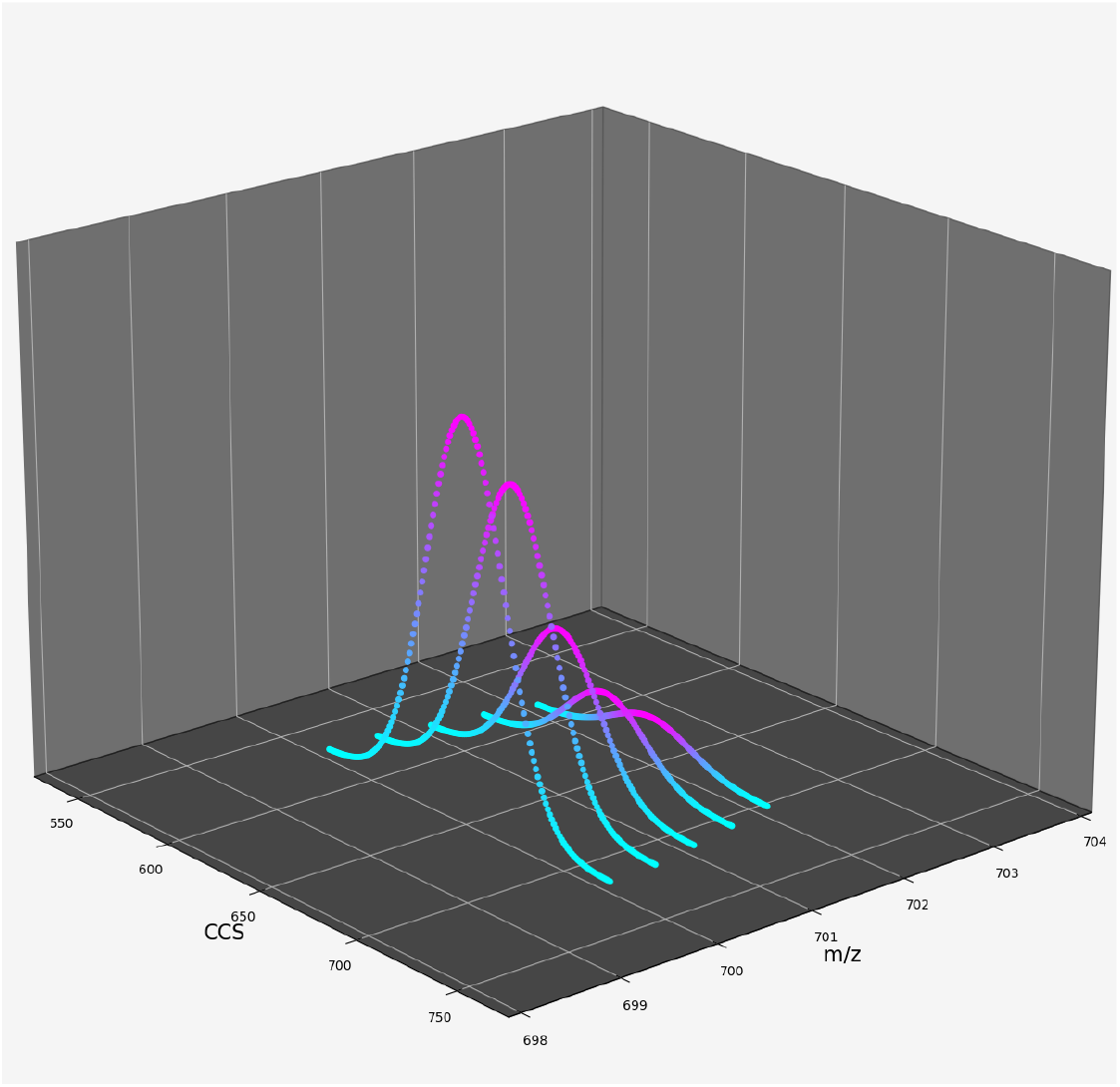
An artificial representation of a peptide’s isotopic peak series in the m/z, CCS, and intensity dimensions at an instant of retention time.

The raw MS1 data space is divided into voxels (volumetric pixels), a small region of 3D m/z, mobility, and retention time space, ranked in descending order by the mean intensity of their constituent raw points. Voxels with fixed dimensions of 0.1 in m/z, 10 scans in mobility, and 5 seconds in retention time are used, where the rationale is to capture a single isotopic peak in the m/z dimension, and to sample the peak in the mobility and retention time dimensions broadly enough to find its apex. A peak encompassed by such a voxel is likely to be a good candidate for a precursor feature’s monoisotopic peak or its base peak. As there are many thousands of voxels to process, processing parallelism is achieved by dividing the m/z dimension into bands of 10 Th through retention time and mobility; a worker is assigned a band in which to process its voxels.

To group the raw points of the candidate peak, the voxel’s intensity-weighted m/z centroid is determined. The peak delta either side of the centroid is determined by calculating three standard deviations based on the instrument’s resolution of 40,000. The raw points within 3*σ* either side of the m/z centroid are grouped as the peak’s m/z dimension. An example voxel, its candidate peak constrained in the m/z dimension, and the raw data in its vicinity are shown in Figure 4.

**Figure 4.**
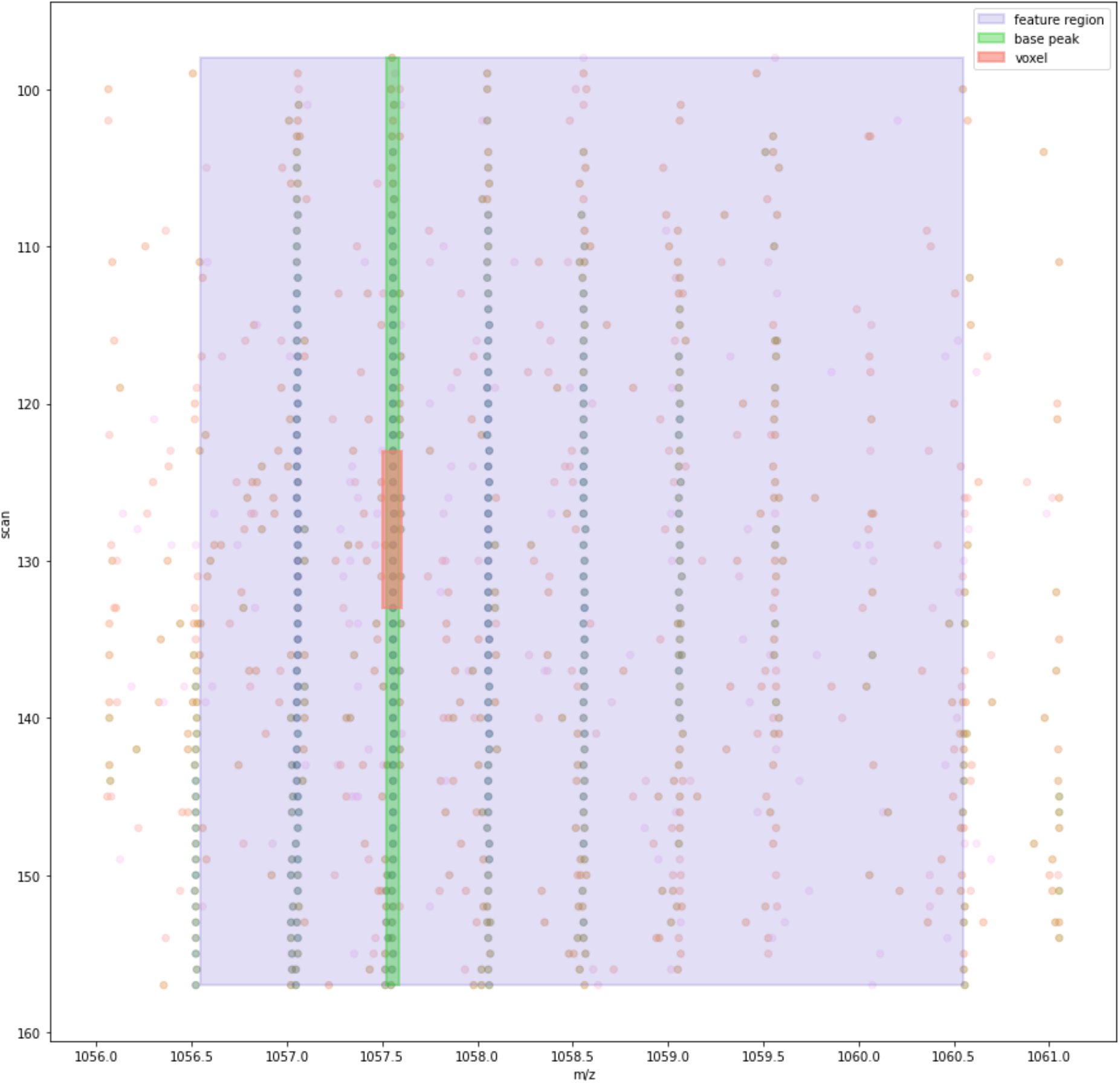
the raw data in the vicinity of an example voxel in its most intense frame

To find the peak’s apex in mobility, we begin by taking points ± 40 scans (a number selected because it is twice the mean peak width in the mobility dimension) from the voxel’s mobility midpoint and flattening the points to the mobility dimension by grouping the points that lie on the same scan and summing their intensity. A smoothing Savgol-Golay filter (13) is applied to facilitate the determination of the peak’s apex and valleys with the peakutils (14) Python package. In instances where we find multiple apexes, we take the apex closest to the voxel’s mobility midpoint. The valley either side of the apex define the peak’s extent in the mobility dimension (Figure 5A).

**Figure 5.**
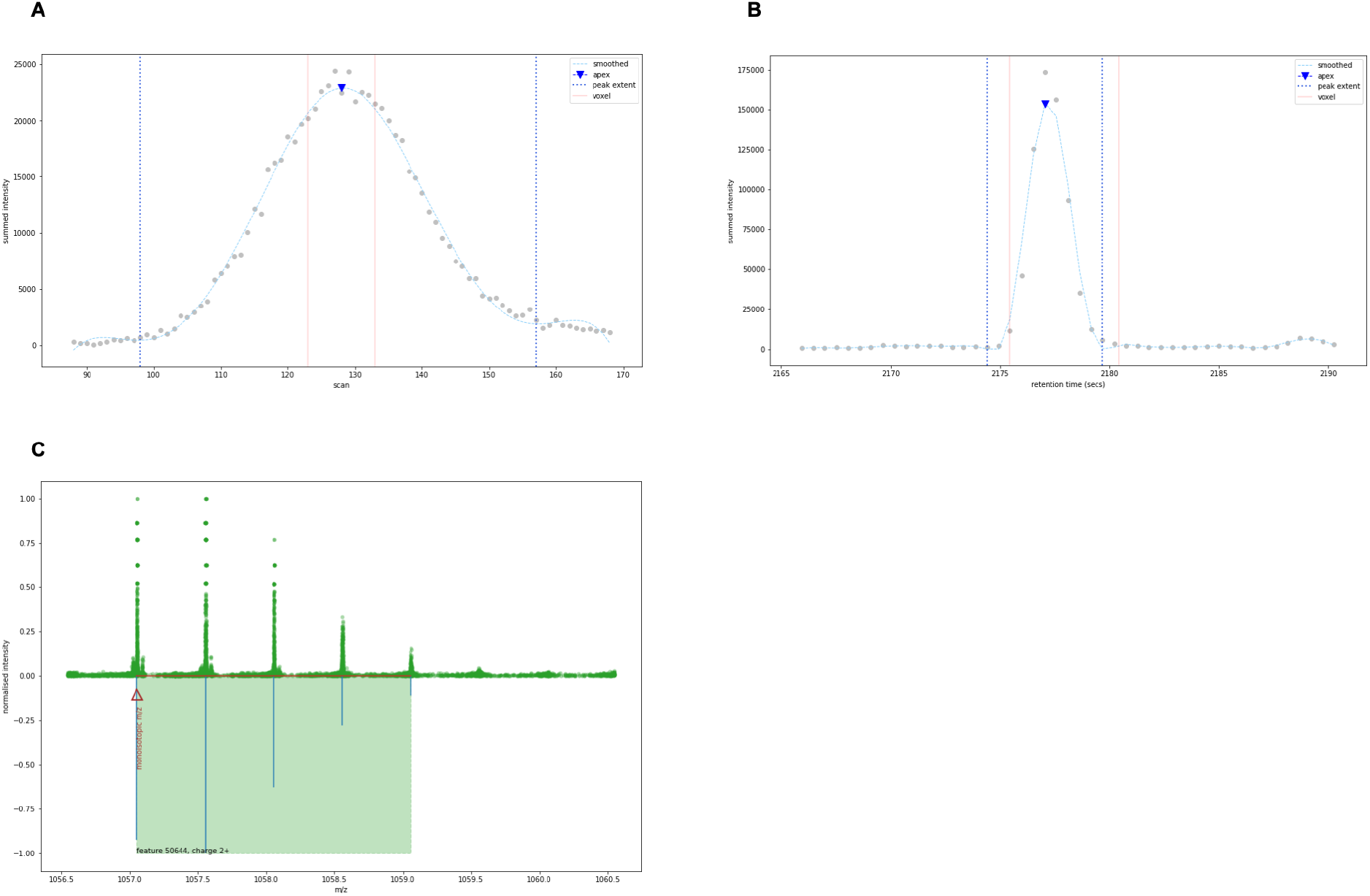
(A) The voxel’s peak flattened to the mobility dimension, with its apex and valleys determing the mobility extent. (B) The same procedeure is used for the retention time dimension. (C) The raw points in the 3D feature region flattened to the m/z dimension, and the isotopic peaks proposed for a charge-2 peptide.

Constraining the raw points to the peak’s m/z and mobility extents, the retention time dimension is extended by twice the mean base peak width in each direction from the voxel’s midpoint. Flattening the points to the retention time dimension by summing the peak’s points that occur in the same frame, the peakutils package is again applied to find the apex closest to the voxel’s midpoint and the valleys on either side. Any points that sit more than 1 second from the main peak are removed to avoid superfluous trailing and leading edges (Figure 5B).

Having defined the bounds of the voxel’s peak in m/z, mobility, and retention time, the region of raw data is expanded in each direction of the m/z dimension to look for the other isotopic peaks in the feature’s series. The lower m/z edge is extended by 0.6 Th (allowing for one isotope for charge-2 plus a little bit more as a margin of error, as the voxel’s peak may be the base peak rather than the monoisotopic peak) and the upper m/z edge by 3 Th (allowing for up to six isotopes for charge-2 plus a little bit more).

The raw points that lie within the 3D feature region are collated, intensity descent as described previously is used to simplify the spectra to 2D (m/z and intensity), and deconvolution is performed with the ms_deisotope Python package (15) to resolve isotopic peaks (Figure 5C).

Each feature that achieves more than the default score is added to the list if it satisfies the requirement of the monoisotopic peak or the base peak matching the voxel’s m/z centroid; this ensures that each feature produced is formed from the voxel’s peak.

Having determined the feature’s monoisotopic m/z, charge state, and isotope envelope, the intensity is calculated for each isotope in the envelope by taking the most intense point in the three frames closest to the retention time apex; three points are used to achieve an averaging effect, to reduce intensity fluctuations that can occur with using single points. If the isotopic peaks include points that were measured when the detector was saturated, their intensities are adjusted with reference to a theoretical model, as we previously reported. The feature’s intensity is calculated by summing the intensity of the first three isotopic peaks. Lastly, voxels that have more than 80% of their intensity comprised of the points in the feature’s isotopic peaks are removed from consideration as the basis of other features.

Voxels are processed in decreasing order of intensity if they have not already been processed in a more intense voxel’s isotopic peaks, continuing until all the voxels above a defined intensity threshold have been processed. The threshold is configurable according to the depth of analysis required; more depth takes more time to process, allowing a trade-off according to the application.

To determine the identifiability of a peptide feature solely from MS1 and in the absence of fragment ion information, inspired by MSTracer (2) we constructed a neural network classifier to predict whether an extracted feature is likely to be identified from MS2 spectra. A training set was created from the identification results from the TFD/E approach we described previously (16), using the features identified with a q-value less than 1% as the ‘identified’ category, and the features detected but not identified as the ‘not identified’ category. The classifier inputs were modified from (2) to suit the additional dimension of mobility and our workflow, as shown in Table 1.

**Table 1 -.** feature attributes used for identifiability classifier inputs

| Attribute | Factor |
| --- | --- |
| deconvolution score | isotopic peak spacing and peak height ratios |
| coelution coefficient | mean cosine similarity of isotopic peaks in retention time |
| mobility coefficient | mean cosine similarity of isotopic peaks in CCS |
| number of isotopes | quality of detection |

The classifier output was the category.

The neural network architecture (Figure 6) was similar to the one used by MSTracer; 3DID’s model was larger in its dimensions as this was determined to improve convergence on on the training data. Defined with Keras (17) and using Tensorflow (18) for the back-end, the dense layers were 200 units wide, batch normalisation was added for each dense layer, and 40% dropout was added to each layer to improve robustness. We randomly selected 80% for training, 10% for validation, and 10% for test.

**Figure 6.**
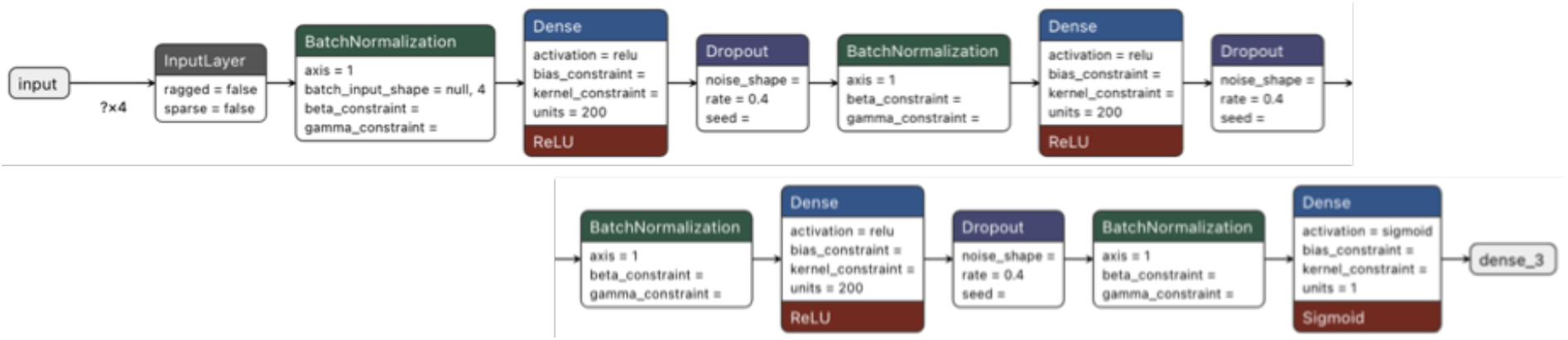
the neural network architecture for the feature identifiability classifier

The model was trained for 4000 epochs with a batch size of 512, which achieved an accuracy of 86% on the test set.

Having been trained on features identified with TFD/E, the model was used to classify the features detected by 3DID. A threshold of 0.2 for identifiability categorisation was selected from examination of the score distribution to filter out only the least likely to be identified and to pass the remainder through to the search engine.

Following the classification step, a de-duplication step was performed to remove features that were inadvertently detected twice. A filter was applied to take the feature with the highest identifiability score when there were other features within 10 ppm m/z, 20 scans in mobility, and 5 seconds in retention time.

### Comparing feature detection with Biosaur

To compare feature detections from 3DID and the latest version of Biosaur (2.0.3), features detected by both tools were matched according to their proximity in m/z, retention time, and CCS. A feature from 3DID and a feature from Biosaur were considered to be the same feature detected by both algorithms if their apexes were within 25 ppm for m/z, 5 seconds in retention time, and 0.05 1/K0 in ion mobility. Default settings were used for Biosaur except for mass_accuracy (10 ppm) and min_intensity (200). This was done for equivalence with 3DID settings as much as possible.

3DID detected 89% of the features detected by Biosaur (13,577), and nearly ten times more (135,218) (Figure 7A). The histogram of feature intensity (Figure 7B) shows the additional features detected by 3DID were mostly of lower abundance than those detected by Biosaur.

**Figure 7.**
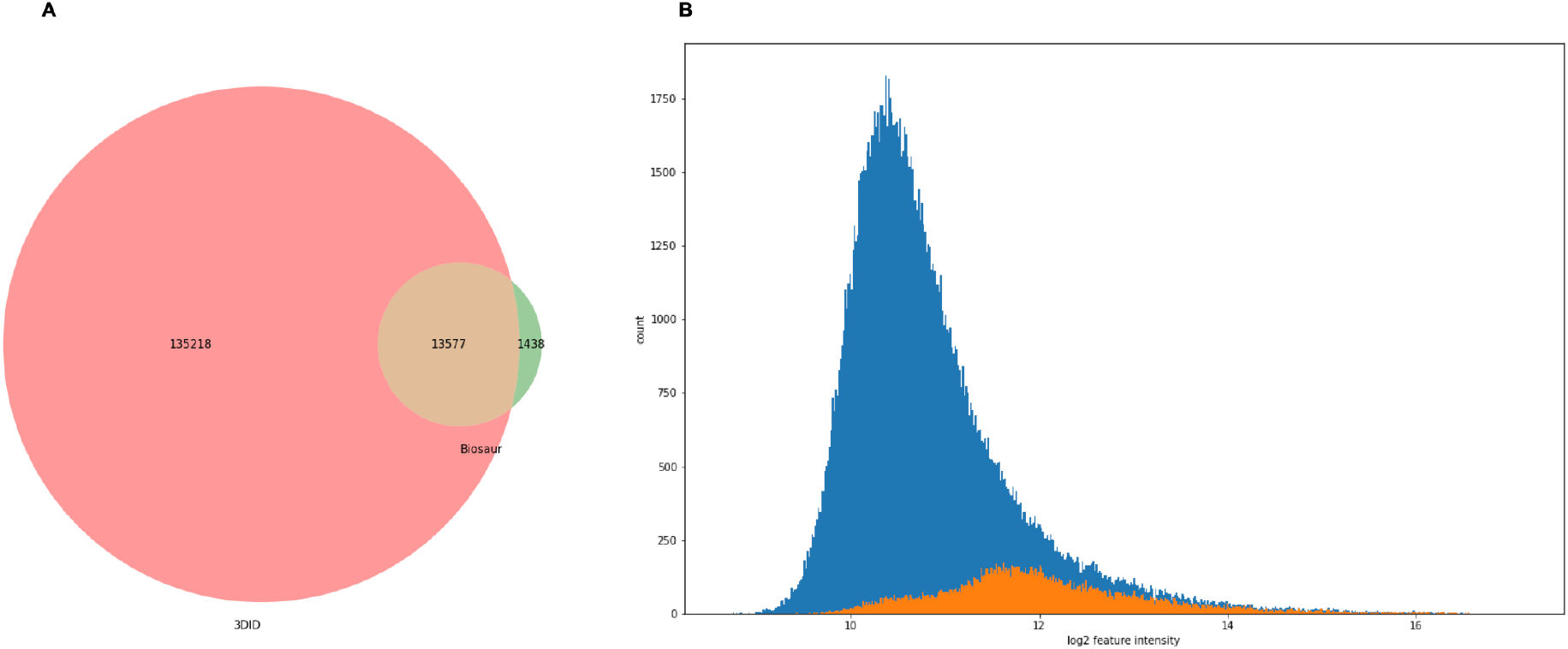
(A) The number of features detected only by 3DID, features detected only by Biosaur, and the features detected by both. (B) The intensity distribution of features only detected by 3DID (blue) and features also detected by Biosaur (orange).

### Comparing identifications with TFD/E and MaxQuant

As MaxQuant only reports detected features that were instrument-selected for fragmentation inside regions recorded as isolation windows, and TFD/E’s feature detection strategy is also based on targeting precursors in isolation windows, a basis for comparison was made by selecting only those features detected by 3DID’s that were inside isolation windows. For a sample of a Yeast/HeLa/E.coli proteome mixture, 156,244 features were detected and classified as identifiable by 3DID at the lowest setting of minimum voxel intensity. A 3DID feature was considered to be located within an isolation window if the apex of its monoisotopic peak was within the bounds of the isolation window’s extent in m/z, retention time, and mobility. We observed that regardless of the minimum voxel intensity setting, the number of features detected in isolation windows was about half the total number of features detected (Figure 8).

**Figure 8.**
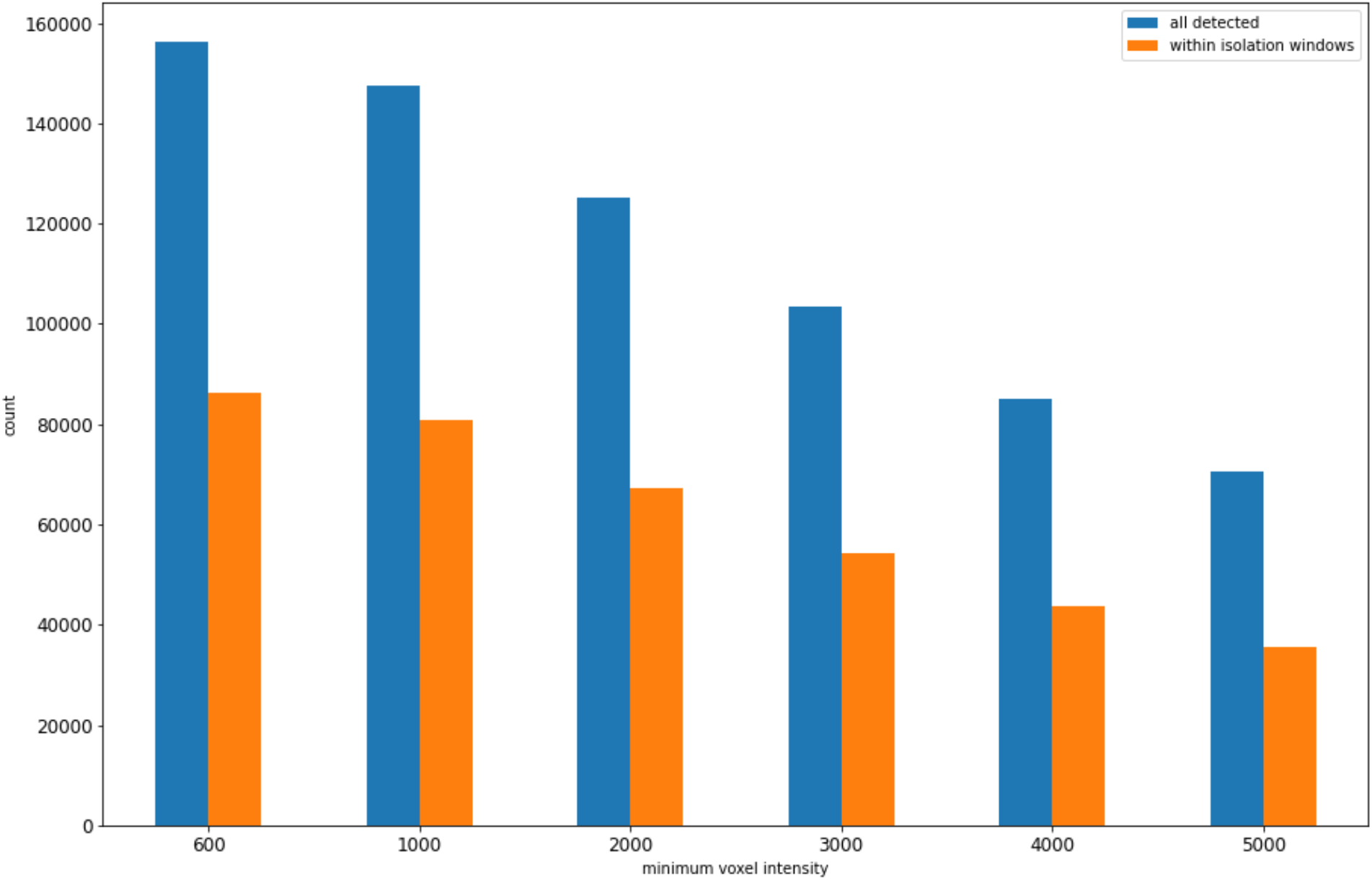
Number of all features detected compared to the number of features within isolation windows, by minimum voxel intensity.

Each feature within an isolation window was associated with the fragment spectra for its window. Using Pyteomics (19), an MGF file (20) was rendered containing an entry for each association, and a search was performed using Comet and Percolator (21) using the same steps from the TFD/E pipeline as previously described: an initial search, mass recalibration, and a more refined search with tighter mass tolerance.

Reducing the minimum voxel intensity at which to end the search for features increases the number of features detected, but the number of identified features does not substantially increase (Figure 9).

**Figure 9.**
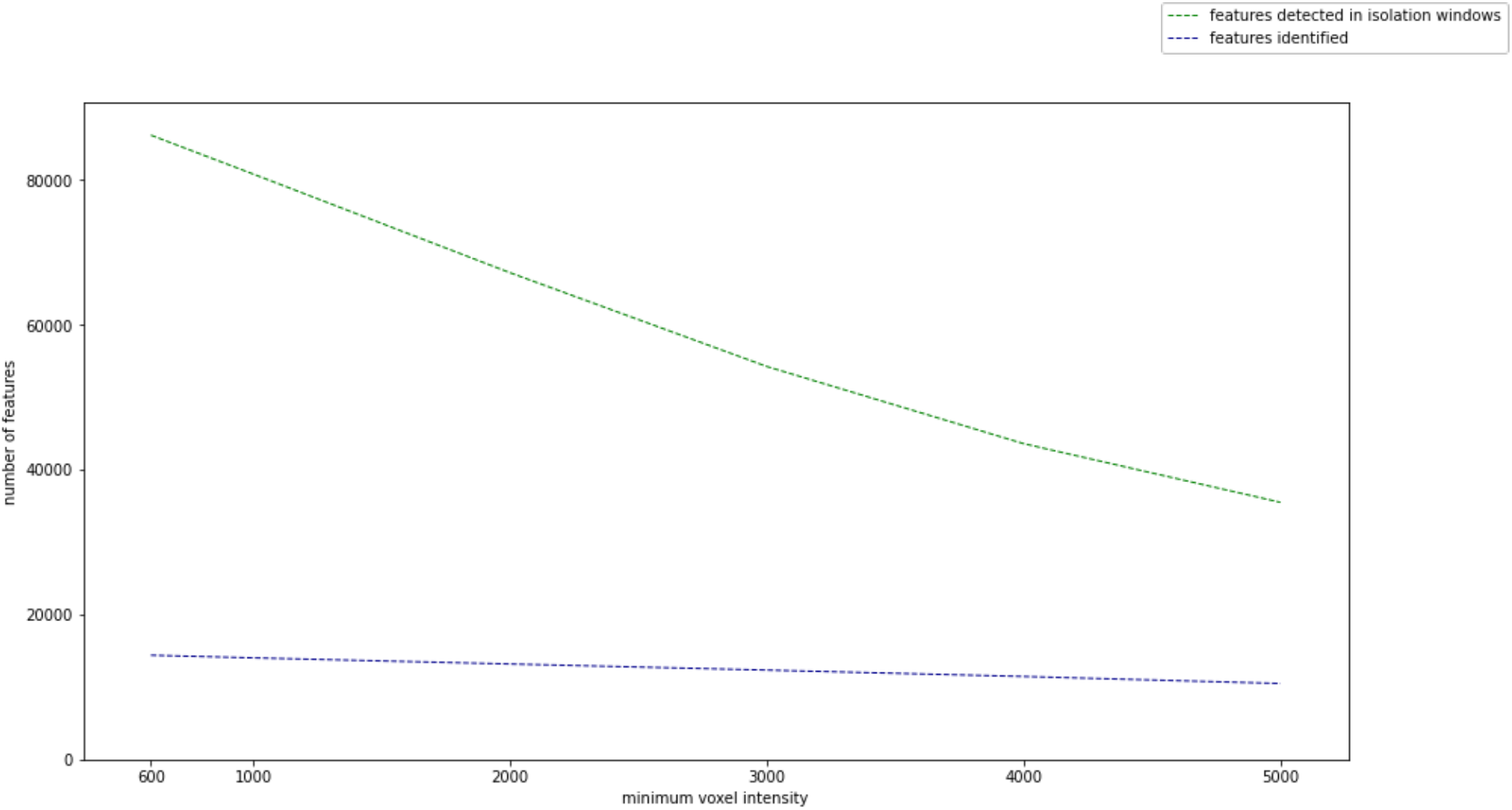
Number of features detected in isolation windows and identified versus minimum voxel intensity.

As the minimum voxel intensity was reduced, the additional features detected were mostly lower intensity, and this would be expected. However, the identified feature intensity distribution did not increase uniformly; many of the additional features detected were not identified (Figure 10). This is likely due to the difficulty of identifying them from the weaker signal of their fragment ions.

**Figure 10.**
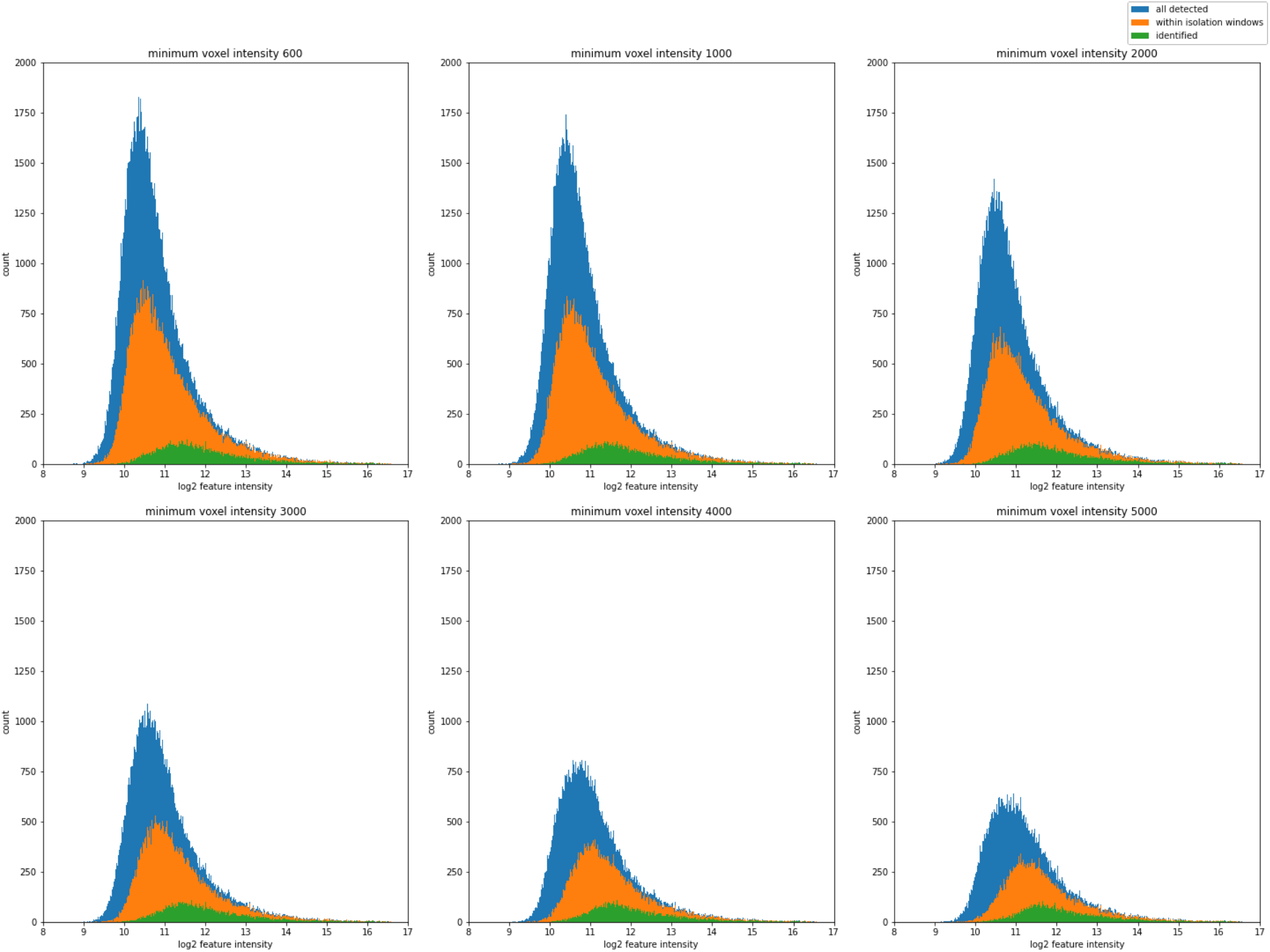
Reducing the minimum voxel intensity increases the number of features detected overall - and within the isolation windows - at an intensity lower than what can be identified with the Comet/Percolator toolchain.

To compare identifications of features detected by MaxQuant while removing the differences between Comet/Percolator and MaxQuant’s integrated search engine Andromeda (8), the features from MaxQuant’s APL peaklist files were rendered as an MGF and searched with Comet and Percolator.

The peptide identifications from features detected by 3DID (within the isolation windows), TFD/E, and MaxQuant had the greatest number of identifications in common at the lower setting for minimum voxel intensity (Figure 11). At the lowest 3DID setting, 7976 unique peptide sequences were identified by all three methods, while a further 1868 peptides were detected by both 3DID and TFD/E.

**Figure 11.**
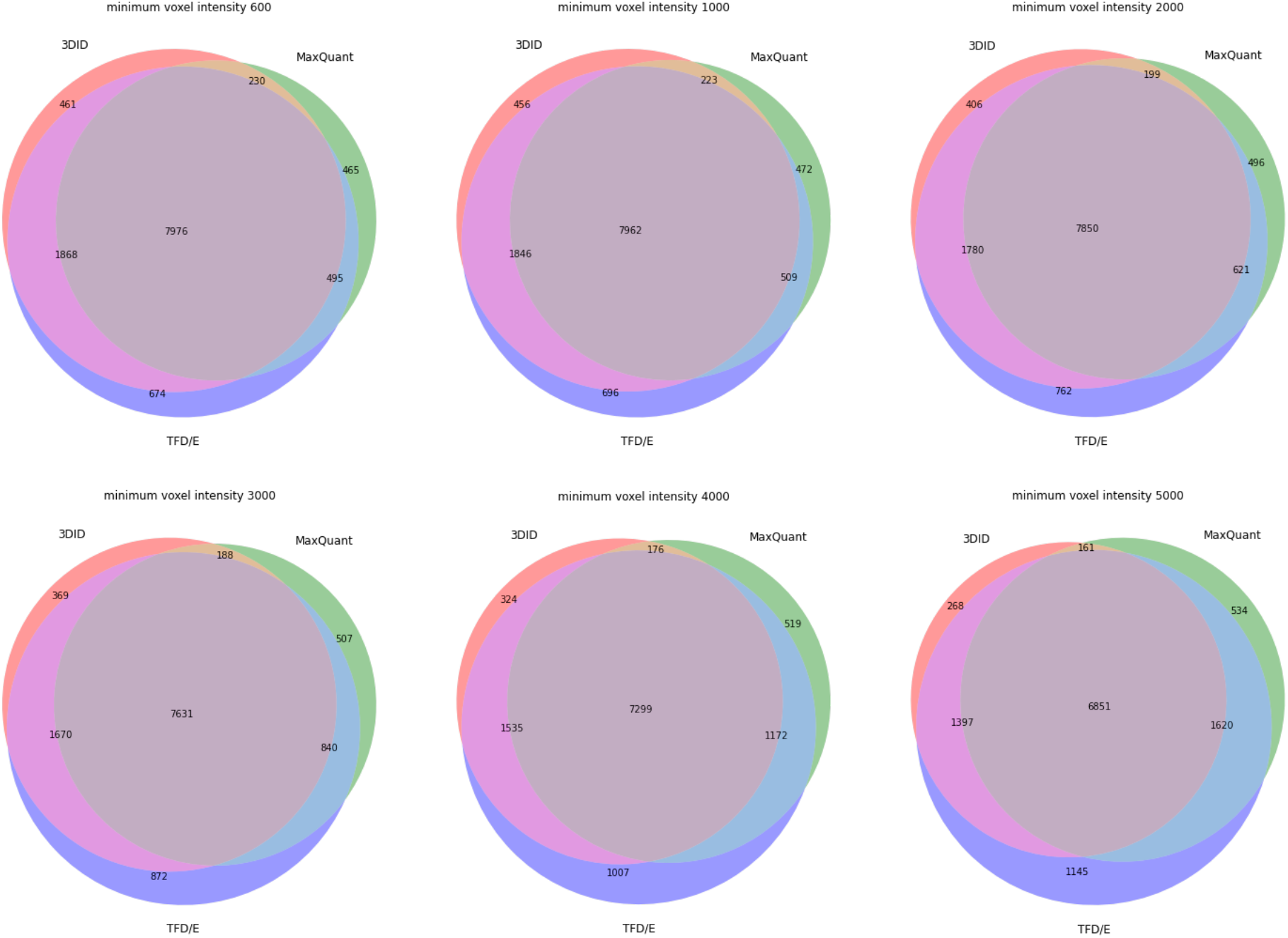
Peptide identifications of the features detected with 3DID and varying minimum voxel intensities 3DID, and the identified features from TFD/E, and MaxQuant.

The increase in identifications from 3DID features as the minimum voxel intensity is lowered does not result in a degradation of mass accuracy; the distribution of mass accuracy is centered around zero within +/- 2ppm and remains consistent with identifications from TFD/E (Figure 12). The bias of mass accuracy for identifications of features detected by MaxQuant is likely due to the absence of a mass recalibration step.

**Figure 12.**
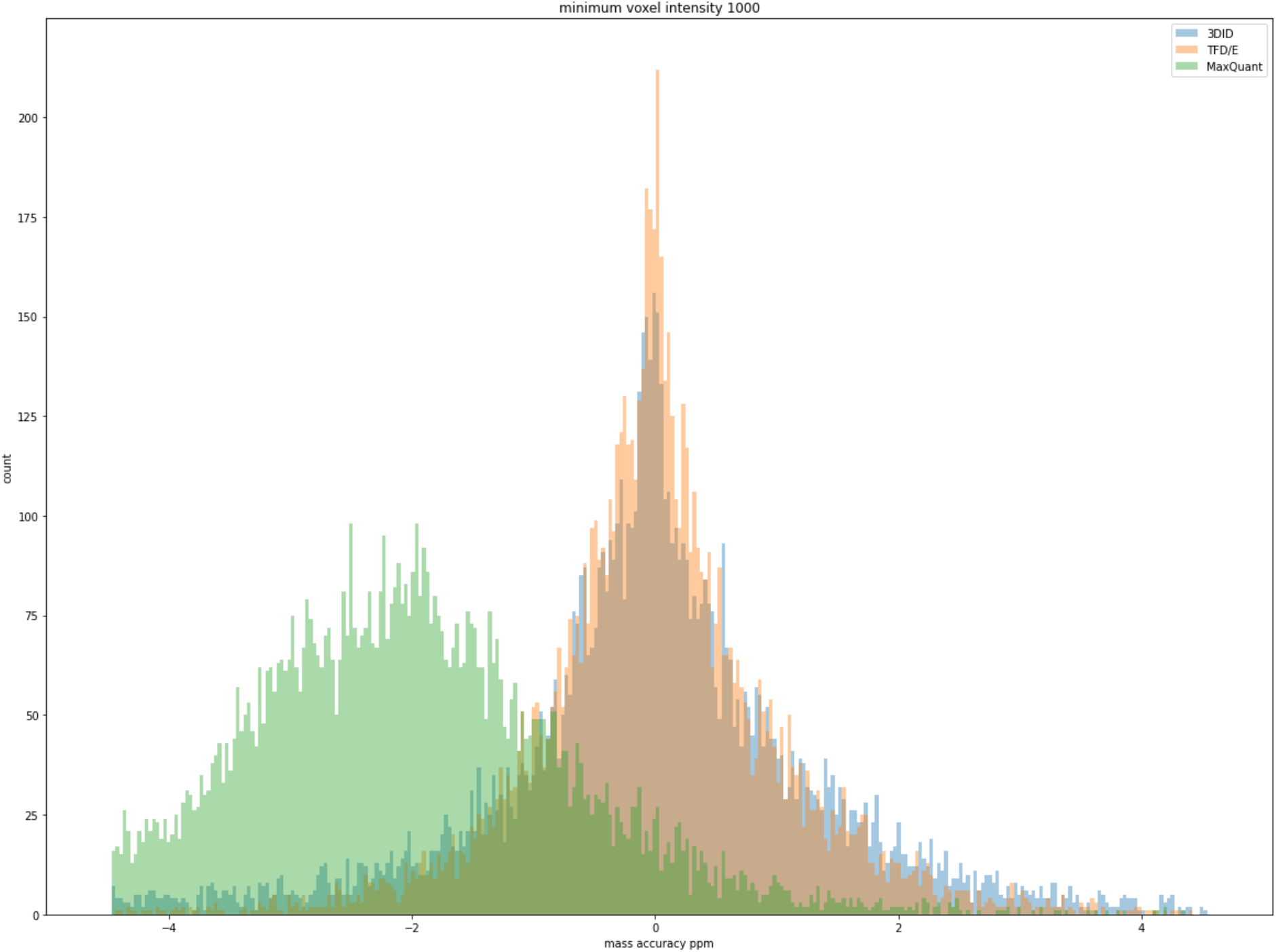
Mass accuracy ppm of identified features detected by 3DID, TFD/E, and MaxQuant.

## 5 Discussion

Here we have presented 3DID, an MS1 feature detector that finds twice as many identifiable features across a broader dynamic range as the DDA analysis approaches of TFD/E and MaxQuant. We have also shown that 3DID detects many more features across a wider dynamic range compared with Biosaur.

The detection of low-abundance peptides is hampered by ambient noise, and the opportunity for false discovery is substantial. However, by imposing a feature quality threshold that considers the feature’s fit with the theoretical tryptic peptide model for its monoisotopic peak and charge state, the alignment of its isotopic peaks in retention time and ion mobility, and the number of isotopes detected, the rate of false discovery is controlled.

3DID’s ability to comprehensively survey precursor ions highlights its potential utility for the characterisation of precursor ions in timsTOF MS1 data.

## 6 Materials and Methods

### 6.1 Data Processing

The YHE211 data set we described previously (16) (available from the ProteomeXchange Consortium via the PRIDE (22) partner repository with the dataset identifier PXD030706 and 10.6019/PXD030706) was processed by 3DID, MaxQuant (2.0.3.0), Biosaur (2.0.3), and TFD/E (1.0).

The raw data was de-noised with the “tims data reduction” tool from Bruker (1.4) using the “denoised proteomics complex” settings.

Default settings were used for MaxQuant. The APL files were converted to MGFs with TFD/E’s generate-MGF-from-MaxQuant-APLs.py.

For Biosaur, the raw data was converted to mzML format with MSConvertGUI from ProteoWizard (3.0) as described in the Biosaur paper (7). Biosaur was run with the following default parameters overridden to match 3DID’s settings as closely as possible: mass accuracy 10, minimum hill length 3, and minimum intensity 200.

Default parameters were used for 3DID and TFD/E.

### 6.2 Software

The software was written in Python 3.8. The key libraries used were Pandas 1.3.1 for data filtering and interface file input/output, AlphaTims 0.3 for loading raw data from the instrument database, scipy 1.6.1 and numpy 1.19.5 for signal processing, ms_deisotope 0.0.22 for spectra deconvolution, and Ray 1.5.2 for parallel processing. The neural network classifier was built with Keras and used TensorFlow 2.5 for the backend. Algorithm prototyping was done in Jupyter notebooks (jupyter-core 4.6.3).

Software validation work was performed on a PC with a 12-core Intel i7 6850K processor and 64 GB of memory running Ubuntu 20.04. An NVIDIA GeForce GTX 1070 GPU was used for the neural network training and inference.

Readers are encouraged to browse the source code in the GitHub repository (DOI 10.5281/zenodo.6513126) for a detailed understanding of the algorithms and implementation approach. The Jupyter notebooks developed to generate the figures in this paper are also available in the repository.

## 7 Acknowledgements

The authors gratefully acknowledge the contributions of Sven Brehmer at Bruker for discussions and guidance about processing timsTOF data.

## Notes

https://doi.org/10.5281/zenodo.6513126

http://dx.doi.org/10.6019/PXD030706

### Competing Interest Statement

The authors have declared no competing interest.

### Summary of Updates

Updated repo URL.

